# High molecular weight DNA isolation method from diverse plant species for use with Oxford Nanopore sequencing

**DOI:** 10.1101/783159

**Authors:** Brieanne Vaillancourt, C. Robin Buell

## Abstract

The ability to generate long reads on the Oxford Nanopore Technologies sequencing platform is dependent on the isolation of high molecular weight DNA free of impurities. For some taxa, this is relatively straightforward; however, for plants, the presence of cell walls and a diverse set of specialized metabolites such as lignin, phenolics, alkaloids, terpenes, and flavonoids present significant challenges in the generation of DNA suitable for production of long reads. Success in generating long read lengths and genome assemblies of plants has been reported using diverse DNA isolation methods, some of which were tailored to the target species and/or required extensive labor. To avoid the need to optimize DNA isolation for each species, we developed a taxa-independent DNA isolation method that is relatively simple and efficient. This method expands on the Oxford Nanopore Technologies high molecular weight genomic DNA protocol from plant leaves and utilizes a conventional cetyl trimethylammonium bromide extraction followed by removal of impurities and short DNA fragments using commercially available kits that yielded robust N50 read lengths and yield on Oxford Nanopore Technologies flow cells.

## Background

The rapid advancement of genomic technologies has enabled a wide range of communities to address long standing questions in medicine, biology, ecology, and evolution. Sequencing technologies have advanced from high throughput Sanger sequencing to ultra-high throughput short read sequences currently dominated by the Illumina platform to third generation long read sequencing platforms such as Pacific Biosciences and more recently, Oxford Nanopore Technologies (ONT). The low infrastructure requirements for ONT has engaged a wide range of users within the broader scientific community and consequently, propelled advancements in not only methodologies but also diversification of use of the ONT platform.

One clear application for ONT sequencing is *de novo* genome assembly in which long reads facilitate generation of long, contiguous assemblies. For generation of long reads on the ONT platform, DNA needs to be high molecular weight, and free of contaminants and nicks. Isolation of high molecular weight DNA is relatively routine for species such as bacteria and mammalian cell culture. However, for plants, the presence of cell walls composed of carbohydrates and lignin as well as a wide range of specialized metabolites including phenolics, terpenes, alkaloids, and flavonoids can impede isolation of high quality and/or high molecular weight DNA. If present during library construction or sequencing on the ONT platform, these impurities can negatively impact yield. One widely adapted method of plant DNA isolation entails inclusion of cetyl trimethylammonium bromide (CTAB) in the extraction buffer which facilitates separation of carbohydrates from DNA (Saghai-Maroof et al., 1984). However, not all impurities are readily removed in CTAB preparations. Furthermore, even limited mechanical activity employed in DNA isolation can result in shearing and nicking of the DNA (Yoo, 2011) which can contribute to yield and/or read length reductions on the ONT platform. Recent reports of *de novo* genome assemblies of plant species that employed ONT utilized extraction of DNA via agarose-embedded nuclei (Deschamps et al., 2018), CTAB extraction followed by a column cleanup (Michael et al., 2018), and custom extraction methods (Belser et al., 2018).

Plants have extreme variation in metabolic diversity across tissues and species as well as genome size not only within a species but also within different aged tissues due to endoreduplication (Del Prete et al., 2019, Laimbeer et al., 2017, Larson-Rabin et al., 2009); both of these can impact the quality and yield of DNA isolation. Indeed, while annual species produce genetically identical true seed from which young, homogenous seedling tissue with limited specialized metabolite composition can be used for DNA isolation, perennials may only produce immature leaf tissue seasonally. As a consequence, mature leaf material or other organs may need to be used for DNA isolation. Furthermore, depending on leaf anatomy and phenology, leaves of some species may have extensive cuticle and/or vascular tissues that are typically enriched in specialized metabolites that can confound isolation of DNA free of contaminants. To avoid generation of a DNA isolation method tailored to each species, we developed a robust species-agnostic and simple protocol for the isolation of pure high molecular weight from diverse plant species which span angiosperm phylogeny, genome size, metabolic diversity, and leaf anatomy and phenology to enable reliable DNA isolation suitable for genome assembly using the ONT platform.

## Methods

### DNA Isolation

DNA was isolated from tissue culture-grown shoots of *Solanum tuberosum* Group Phureja clone DM1-3 516 R44 (potato) and leaves of *Lavandula angustifolia* (lavender) using the Workman et. al (2018) protocol followed by the Nanobind Plant Nuclei Big DNA Kit (Circulomics, Baltimore, MD, Cat # NB-900-801-01). *Lavandula angustifolia* DNA was also isolated using a CTAB method (Doyle & Doyle, 1987). DNA was isolated from *Arabidopsis thaliana* (Arabidopsis), *Ipomoea batatas* (sweetpotato), *L. angustifolia, Nepeta cataria* (catnip), and *Nepeta mussinii* (catmint) (Table 1) following the protocol provided by Oxford Nanopore Technologies, “High molecular weight gDNA extraction from plant leaves” downloaded from the ONT Community in February, 2019 (CTAB-Genomic-tip). This protocol reports on isolation of DNA from spinach leaves using a CTAB buffer followed by a Qiagen Genomic-tip cleanup and size selection using AMPpure XP beads. Metrics reported were DNA yield, DNA fragment size, DNA purity, as well as yield (8+ Gb), read length distribution, and sequenced base quality on an ONT flow cell. In our adaptation of this protocol, young leaves were harvested, flash frozen in liquid nitrogen, and ground to a very fine powder in liquid nitrogen. Approximately one gram of ground tissue was transferred to 20 ml of Carlson buffer (100 mM Tris-HCl pH 9.5, 2% CTAB, 1.4 M NaCl, 1% PEG 8000, 20 mM EDTA, 0.25% β-mercaptoethanol (v/v)) prewarmed to 65 °C. To ensure complete removal of RNA from the sample, 40 μl of RNase A (100 mg/ml) was added to the sample, incubated at 65 °C for thirty minutes; subsequently, an additional 40 μl of RNase A was added and incubated another 30 minutes. After cooling to room temperature, one volume of chloroform was added, vortexed, and spun at 5,500 g for 10 minutes at 4 °C. The aqueous layer was then mixed thoroughly with 0.7 volumes of isopropanol, incubated at −80 °C for 15 minutes, and then centrifuged at 5,500 g for 30 minutes at 4 °C; the pellet was dissolved at 50 °C for 15 minutes in 19 ml of Buffer G2 (Qiagen, Germantown, MD, Cat #1014636).

**Table 1:** List of species used in this study with taxonomic clade and genome size.

| Species | Taxonomic Clade | Genome Size | Reference |
| --- | --- | --- | --- |
| *Arabidopsis thaliana* | Brassicaceae | 157 Mb | Bennett et al. 2003 |
| *Ipomoea batatas* | Convolvulaceae | 4.4 Gb | Ozias-Akins & Jarret 1994 |
| *Lavandula angustifolia* | Lamiaceae | 968 Mb^ | This study |
| *Nepeta cataria* | Lamiaceae | 733 Mb^ | This study |
| *Nepeta mussinii* | Lamiaceae | 303 Mb^ | This study |
| *Solanum tuberosum* Group<br>Phureja | Solanaceae | 844 Mb | Potato Genome<br>Sequencing Consortium<br>2011 |

^Size estimated by flow cytometry.

After equilibrating the Qiagen Genomic-tip 500/G column (Qiagen, Germantown, MD, Cat #10262) with 10 ml of Buffer QBT (750mM NaCl • 50 mM MOPS pH 7.0, 15% isopropanol (v/v), 0.15 % Triton X-100 (v/v)), the resuspended DNA was applied to the column, washed twice with 20 ml of Buffer QC (1M NaCl, 50 mM MOPs pH 7.0, 15% isopropanol (v/v)), and eluted with 15 ml of Buffer QF (1.25 M NaCl, 50 mM Tris-Cl pH 8.5, 15% isopropanol (v/v)) prewarmed to 55 °C. The DNA was precipitated with 0.7 volumes of isopropanol at room temperature for 15 minutes and centrifuged at 5,500 g for 30 minutes at 4 °C. The pellet was then washed with 2 ml of 70% ethanol, dried, and eluted in Buffer EB (10 mM Tris-Cl, pH 8.5) overnight at room temperature; DNA was stored at 4 °C long term.

### Removal of Impurities using Amicon Buffer Exchange

An Amicon buffer exchange was performed with the CTAB-Genomic-tip isolated DNA, starting with a pre-rinse of the Amicon Ultra-0.5 Centrifugal Filter Unit (Millipore, Burlington, MA, Cat #UFC510008) with 100 μl of nuclease free water spun at 14,000 g for three minutes. Any residual water that did not flow through the filter was discarded. The DNA sample was applied to the filter, volume brought to 500 μl with Buffer EB, and spun at 14,000 g for ten minutes. The filtrate was discarded, 500 μl of Buffer EB applied to the filter and spun at 14,000 g for a time dependent on the desired final volume. The filter was then inverted into a new tube and spun at 1,000 g for two minutes; all spins were performed with the cap hinge towards the center of the centrifuge rotor. DNA was stored at 4 °C long term.

### Removal of Short Fragments

To enhance recovery of long DNA fragments, the Short Read Eliminator Kit (Circulomics, Baltimore, MD, Cat #SS-100-101-01) v1.0 protocol from February, 2019 was used; it is recommended to follow the protocol most recently released by Circulomics. The maximum input of 9 μg of DNA in 60 μl (150 ng/μl) was used, diluting with Buffer EB in an Eppendorf DNA LoBind tube (Eppendorf, Hauppauge, NY, Cat #13-698-791). A total of 60 μl of Buffer SRE was added, mixed by gently flicking the tube, and centrifuged at 10,000 g for 30 minutes at room temperature. The supernatant was removed, and the pellet washed twice with 200 μl of 70% ethanol by centrifuging at 10,000 g for two minutes. The DNA pellet was then dissolved in 50 μl of Buffer EB by incubating at 50 °C for one hour. To make sure the DNA was fully resuspended the tube was gently tapped. DNA was stored at 4 °C until library preparation. Libraries should be prepared as soon as possible after the Short Read Eliminator protocol as degradation of DNA can occur during storage resulting in shorter fragments of DNA. Storing the sample for fewer than two days was found to result in fewer short reads during sequencing.

### Oxford Nanopore Technologies Library Preparation, Sequencing, and Base Calling

DNA libraries were prepared using the Oxford Nanopore Technologies SQK-LSK109 kit with the following modifications. DNA repair and end-prep (New England BioLabs, Ipswich, MA, Cat #E7546 and Cat #M6630) were performed with 3 μg DNA, in two separate tubes (1.5 μg in each tube) in a total reaction volume of 60 μl each, incubated at 20 °C for 60-90 minutes, and 60 °C for 60-90 minutes. The two DNA repair and end-prep reactions were combined and cleaned with 120 μl of Agencourt AMPure XP beads (Beckman Coulter, Brea, CA, Cat #A63880) with an incubation time ranging from ten to 20 minutes and an elution time of five to ten minutes. Ligation was performed at room temperature in the dark for two hours. The ligation reaction was cleaned using Agencourt AMPure XP beads with an incubation time of ten to 20 minutes and an elution time of 15 to 25 minutes at room temperature or 37 °C (recommended). The entire library was loaded onto the flow cell. We investigated several HulaMixer (Invitrogen, Carlsbad, CA, Cat #15920D) settings for both bead incubation steps; the final preferred setting was orbital rotation = 7, orbital rotation time = OFF, reciprocal motion turning angle = 90 °, reciprocal motion time = 15, vibrating motion turning angle = 1 °, vibrating motion = OFF.

Sequencing was performed on an Oxford Nanopore Technologies (Oxford, UK) MinION (MIN-101B) with FLO-MIN106 Rev D flow cells using an Apple Macintosh computer or MinIT (MNT-001). There were several versions of MinKNOW (https://community.nanoporetech.com/downloads) released during this study, the following versions were the most up-to-date version of MinKNOW available when sequencing was performed: v3.1.19, 3.1.20, 3.3.2, 3.3.3. MinKNOW parameters were set to default except for allowing a maximum run time of 96 hours.

All samples in Table S1 as well as *L. angustifolia* in Table S2 were base called using Guppy v3.1.5 (https://community.nanoporetech.com/downloads) in fast mode. The following parameters were set: config dna_r9.4.1_450bps_fast.cfg, trim_strategy dna, qscore_filtering, calib_detect, and calib_reference lambda_3.6kb.fasta, all other parameters were left at default. *S. tuberosum* Group Phureja data was based called using Guppy (v3.2.2+9fe0a78) with the options --qscore_filtering, -- trim_strategy dna, and --calib_detect.

Detailed methods and statistics for the library preparation, sequencing, and base calling of the DNA prepared using the CTAB-Genomic-tip method with and without the Amicon cleanup and Short Read Eliminator are provided in Table S1. Throughout the course of this experiment and in developing this protocol there were numerous MinKNOW, Guppy, flow cell, and computational updates. Details for DNA isolation method, library preparation, and sequencing for the other DNA isolation methods are available in Table S2.

## Results & Discussion

For the doubled monoploid *S. tuberosum* Group Phureja clone DM1-3 516 R44 with a genome size of 844 Mb (The Potato Genome Sequencing Consortium, 2011), we successfully performed a nuclei extraction following the Workman et. al (2018) protocol. Nuclei were then used as input into the Nanobind Plant Nuclei Big DNA Kit to isolate DNA and sequence on the ONT platform. In total, we sequenced *S. tuberosum* Group Phureja clone DM1-3 516 R44 on six flow cells yielding an average N50 of 42.5 kb with an average yield per flow cell of 7.0 Gb (base called passed reads); one library was run on a FLO-MIN106 and the rest on FLO-MIN106 Rev D flow cells (Table S2). We also utilized the Nanobind kit to isolate DNA from the 968 Mb genome sized *L. angustifolia*, a species well known for its production of secondary metabolites including volatile terpenoids associated with fragrance. While we were able to successfully isolate high molecular weight DNA from leaves with an average flow cell yield of 6.17 Gb and average N50 of 30.0 kb when sequenced on three FLO-MIN106 and two FLO-MIN106 Rev D flow cells (Table S2), in subsequent efforts, we were unable to isolate any DNA from *L. angustifolia* using nuclei isolation and the Nanobind kit. Alternative extraction of DNA from *L. angustifolia* using a standard CTAB extraction buffer with a subsequent buffer exchange and the Circulomics Short Read Eliminator that enriches DNA preparations for fragments greater than 25 Kb were also unsuccessful in generating adequate yield on the flow cell (1.20 and 1.38 Gb; Table S2). While the genome of Arabidopsis has been available since 2000 (Arabidopsis Genome Initiative, 2000), generation of whole genome assemblies of additional accessions has value in determining allelic and structural variation among accessions (Pucker et al., 2019, Zapata et al., 2016). We attempted isolation of DNA from the small sized genome of *A. thaliana* (157 Mb) using the nuclei isolation and Nanobind Plant Nuclei Big DNA Kit; surprisingly, we were unable to recover sufficient DNA from Arabidopsis (data not shown).

For the ONT platform to be efficient, we require a high molecular weight DNA isolation protocol that is not only efficient with respect to labor and reagent/material costs but also effective and consistent with diverse species that span a range of genome sizes, have diverse specialized metabolite composition, and produce leaves with differing phenology. We tested the Oxford Nanopore Technologies, “High molecular weight gDNA extraction from plant leaves” method (CTAB-Genomic-tip) with *L. angustifolia* and *N. mussinii* generating N50 read lengths of 22.6 kb and 20.7 kb and yields of 12.10 Gb and 5.54 Gb, respectively (Table 2; Table S1). As both of these species are known for diverse specialized metabolite production, we explored addition of an Amicon buffer exchange following DNA isolation. This resulted in increased N50 read lengths to 25.5 kb and 25.0 kb for *L. angustifolia* and *N. mussinii*, respectively, with an increased yield in *N. mussinii* yet lower in *L. angustifolia* (Table 2; Table S1). We also explored the inclusion of the Circulomics Short Read Eliminator. As shown in Table 2, the Short Read Eliminator increased N50 read length for both species with and without the Amicon buffer exchange.

**Table 2:**
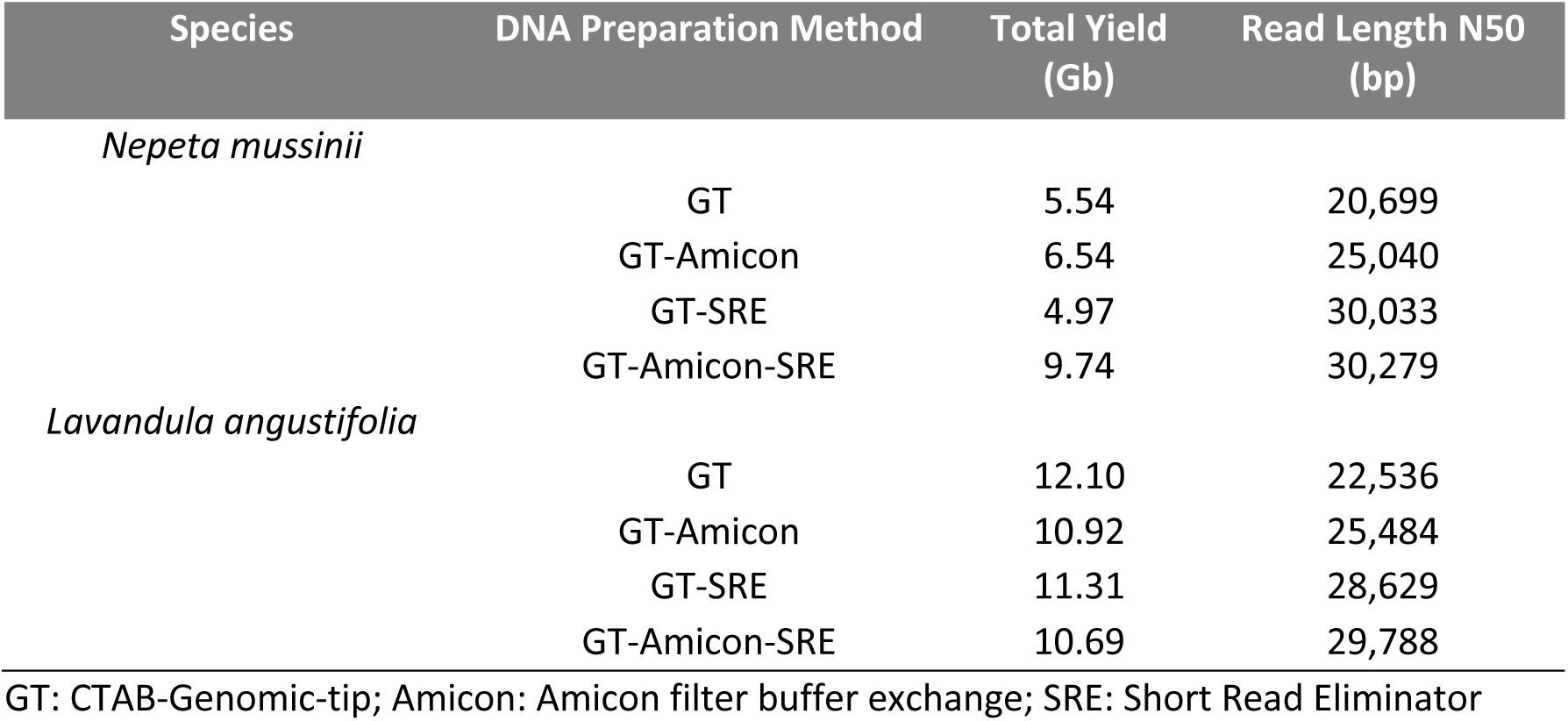
Variation in sequencing metrics with alternative post-isolation preparation methods. Total yield and read length are passed reads after base calling with calibration reads removed.

We then explored the combination of CTAB-Genomic-tip followed by both a buffer exchange with an Amicon filter and size selection with the Short Read Eliminator with three additional species: *N. cataria*, a tetraploid with a 733 Mb genome which is a producer of diverse secondary metabolites, *A. thaliana*, an annual herbaceous species, and *I. batatas*, a hexaploid with a genome size of 4.4 Gb with waxy leaves. For each species, we were able to generate robust N50 read lengths and reasonable yields (Figure 1; Table S1).

**Figure 1:**
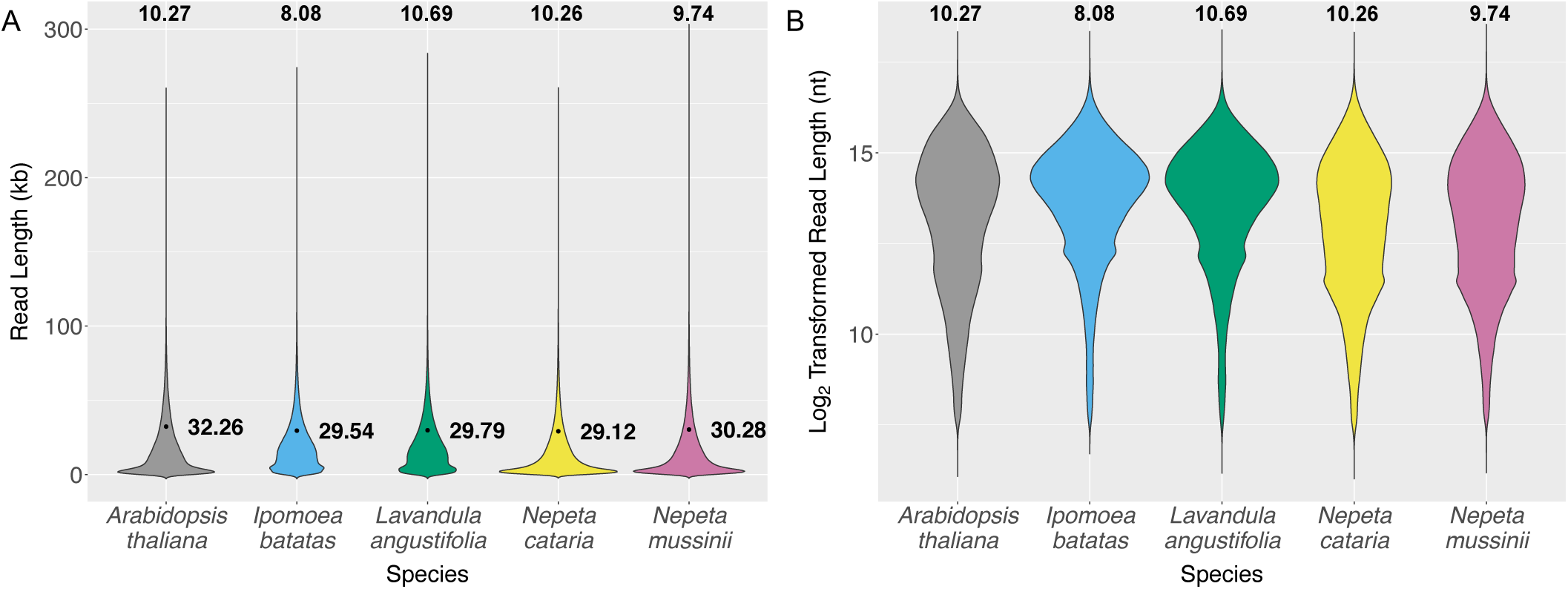
Read length distributions of five species with DNA extracted using the CTAB-Genomic-tip method followed by an Amicon buffer exchange and Short Read Eliminator size selection. Only passed reads excluding calibration reads are presented. **A**. Violin plot of read lengths with the N50 read length to the right of each plot and total yield in gigabases at the top of each plot. **B**. Violin plot of log_2_ transformed read lengths with the total yield in gigabases at the top of each plot.

## Conclusion

With a focus on obtaining long reads and >50X coverage of the genome, two criteria for robust *de novo* genome assemblies, we have found that we are able to obtain sufficient read length distribution and yield by expanding on the ONT CTAB-Genomic-tip protocol by inclusion of an Amicon buffer exchange and size selection via the Circulomics Short Read Eliminator (Figure 2). Although other isolation methods have been successfully used for generating ONT long reads, including within our laboratory, we were able to successfully isolate DNA from five taxa that represent a diverse metabolism, genome sizes, and leaf types using a straightforward protocol without the need for species-specific optimization.

**Figure 2:**
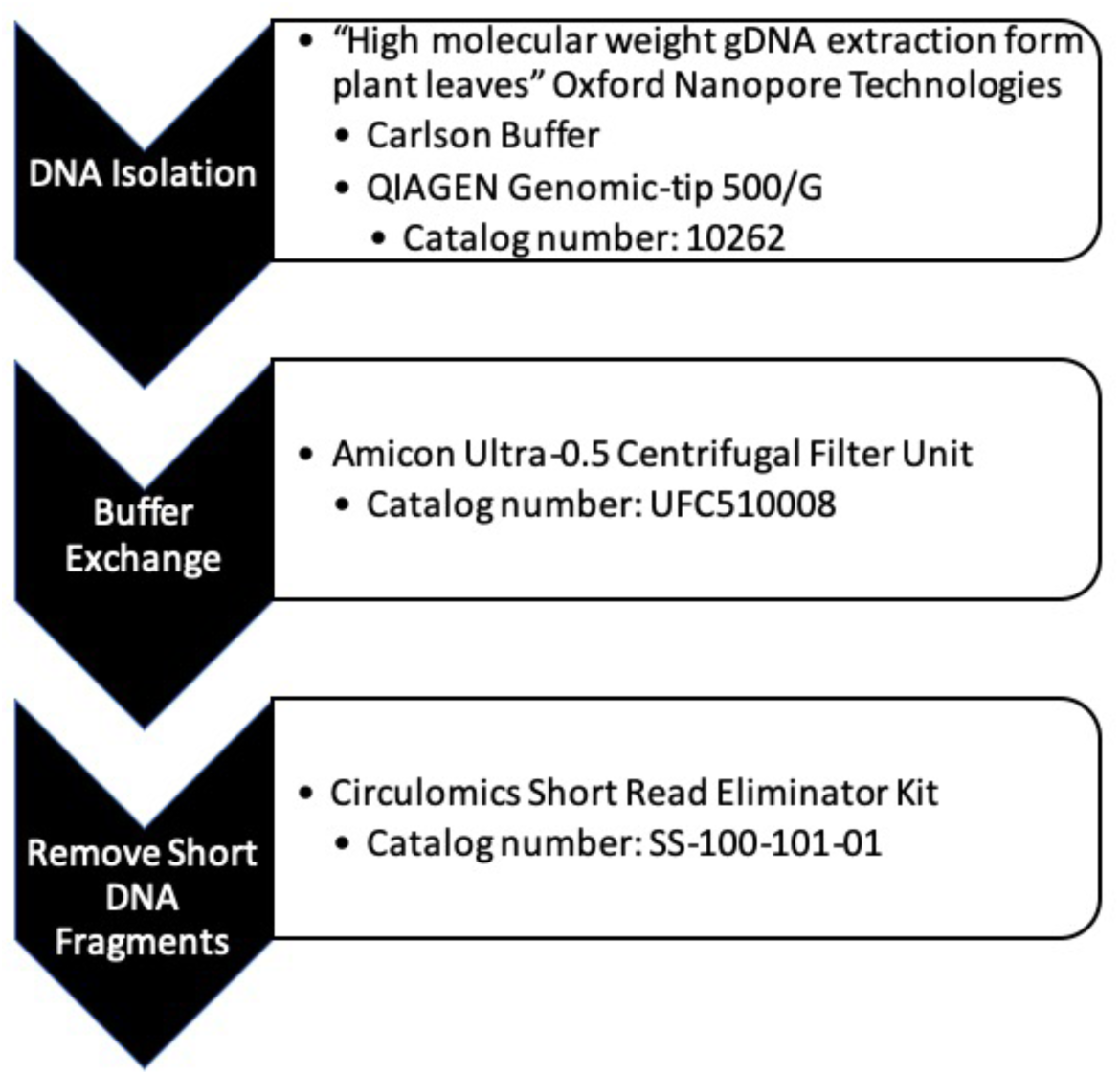
Method for high molecular weight DNA isolation for Oxford Nanopore Technologies sequencing.

## List of Abbreviations

CTAB: cetyl trimethylammonium bromide
ONT: Oxford Nanopore Technologies

## Authors’ contribution

BV and CRB conceived the study. BV performed the experiments. BV and CRB wrote the manuscript. Both authors approved the final manuscript.

## Acknowledgements

We thank the members of the Buell lab and Kevin Childs for robust discussions on ONT. We thank John Hamilton for work on *S. tuberosum*.

